# Influence of perinatal ampicillin exposure on maternal fecal microbial and metabolic profiles

**DOI:** 10.1101/2025.06.30.662372

**Authors:** Simone Zuffa, Sydney P. Thomas, Ipsita Mohanty, Yasin El Abiead, Victoria Deleray, Kine Eide Kvitne, Armin Kousha, Emi Suzuki, Chih Ming Tsai, Griffith Nguyen, Benjamin Ho, George Y. Liu, Victor Nizet, Pieter C. Dorrestein, Fatemeh Askarian, Shirley M. Tsunoda

**Author notes:** Corresponding authors ( for animal work, for metabolome and microbiome). equally contributed (Simone Zuffa, Sydney P. Thomas).

## Abstract

Indirect exposure to antibiotics during early life, via maternal intrapartum antibiotic prophylaxis (IAP) or postpartum maternal antibiotic usage, is increasingly common and has been epidemiologically linked to altered growth and immune developmental trajectories in offspring. Nevertheless, the underlying mechanisms remain poorly understood. Here, we explored the effects of antepartum and postpartum maternal ampicillin administration on the dams’ fecal microbiome and metabolic profiles *in vivo*. Ampicillin caused a reproducible depletion of beneficial bacterial species belonging to the *Muribaculaceae* family, including *Muribaculum intestinale* and *Duncaniella dubosii*, and led to cohort-dependent enrichments of *Enterococcus* and *Prevotella* species. These microbial alterations were accompanied by substantial metabolic remodeling, characterized by elevated fecal acylcarnitines and dysregulation of the bile acids profile. Intriguingly, we identified two previously uncharacterized trihydroxylated bile acids conjugated to a hexose moiety, which appeared to be associated with antibiotic exposure across public metabolomics repositories. These alterations in the fecal maternal microbiome and metabolome coincided with increased weight gain in offspring, suggesting a possible role for maternal antibiotic exposure in shaping early developmental trajectories. Further studies are warranted to elucidate the long-term implications of these changes in infant health.

**IMPORTANCE:** Perinatal antibiotic administration is a critical intervention to reduce maternal and neonatal infections, including early-onset group B *Streptococcus* (GBS) disease, a major cause of neonatal mortality. Nevertheless, mounting evidence suggests that the use of broad-spectrum antibiotics during the perinatal period in mothers can affect infant gut microbiome development, with potential consequences for immune maturation and early development. Understanding how maternal antibiotic exposure affects the gut microbiome and metabolome is essential for uncovering the potential pathways by which maternal intervention may influence offspring outcomes and for guiding strategies that balance infection control with long-term infant health.

## INTRODUCTION

Birth is a defining moment in infant development. During and after delivery, microorganisms from the mother and surrounding environment colonize the newborn (1, 2) and initiate molecular interactions that are fundamental for immune maturation (3, 4), neurodevelopment (5–7), and numerous other biological processes (8, 9). Disruptions to these early gut microbial communities, particularly via external factors such as antibiotic exposure (10–12), have been associated with increased risks for several later-life conditions, such as obesity (13, 14), type 1 diabetes (15), atopic diseases (16, 17), and neurodevelopmental disorders (18, 19). Despite growing awareness of these risks, the impact of indirect exposure to antibiotics via maternal administration remains poorly understood. Prophylactic antibiotic treatment before or during labor is commonly used to reduce complications and infections in both mothers and neonates across high-, middle-, and low-income countries (20, 21). As such, the consequences of these interventions on the infant gut microbiome requires deeper investigation.

Ampicillin, a broad-spectrum β-lactam antibiotic, is commonly administered intravenously to at-risk mothers as intrapartum antibiotic prophylaxis (IAP) to prevent early-onset group B *Streptococcus* (GBS) disease in newborns (22). Approximately 30% of pregnant women test positive for GBS during routine screening in late pregnancy (23) and 1% of the colonized newborns will develop invasive GBS infection (24). Although IAP reduces the risk of early-onset disease (EOD), GBS remains a leading cause of neonatal sepsis and meningitis, with a mortality rate of ∼5% among affected infants (25). EOD survivors are also at greater risk of neurodevelopmental disorders including cerebral palsy, blindness, deafness, and cognitive delay (25). Importantly, IAP does not reduce the incidence of late-onset GBS disease (26), and may facilitate the transmission of antimicrobial-resistant (AMR) bacteria (27). Furthermore, IAP has been shown to disrupt beneficial microbial colonization in neonates, impairing early-life microbiota assembly (28, 29). These observations underscore the need to better understand how perinatal antibiotic administration, particularly the timing of exposure, modulates the maternal gut microbiome and metabolome composition, and its downstream influence on offspring development.

In this study, we investigated the *in vivo* effects of maternal exposure to the broad spectrum antibiotic ampicillin, administered either antepartum or postpartum, on maternal gut microbial and metabolic profiles via shotgun metagenomics sequencing and untargeted liquid chromatography coupled with tandem mass spectrometry (LC-MS/MS) metabolomics. Notably, we observed significant alterations not only in bacterial taxa but also in carnitine and bile acid profiles, metabolites known to play a critical role in energy homeostasis and immune regulation (30, 31), with the potential to influence offspring physiology through maternal transfer. Although both antepartum and postpartum ampicillin exposure were associated with increased offspring weight, the antepartum treatment resulted in more persistent alterations to the maternal fecal metabolome. These findings highlight that the timing of perinatal ampicillin exposure can differentially shape fecal maternal microbiota and metabolome, with implications for neonatal development. Our results emphasize the need to critically evaluate the necessity, spectrum, and timing of prophylactic antibiotic administration surrounding childbirth.

## MATERIAL AND METHODS

### Animal Study and Sample Collection

Wildtype female timed pregnant C57BL/6 mice (strain#000664) were either obtained from the Jackson laboratory at gestational day (GD) 12 or 13 or manually set up for mating for 24 hours using 10- to 12-week-old male and female mice (1:1). Following confirmation of pregnancy on GD14, determined by abdominal palpation and visible enlargement of the abdomen, the animals were randomized into the designated experimental treatments and assigned to one of two cohorts: the Antepartum cohort or the Postpartum cohort. In the Antepartum cohort, animals were administered 150 mg/kg of ampicillin (AMP) or phosphate-buffered saline (PBS) via retro-orbital injection on GD17 and GD18, with delivery predicted on GD19. In the Postpartum cohort, animals received the same retro-orbital injections of AMP or PBS on postnatal days 2 and 3 (PND2 and PND3). Animals were housed in filter-top cages and had *ad libitum* access to standard chow diet and water. Ambient temperature was maintained at 20-22 °C, humidity at 30-70%, and a 12 h light/12 h dark cycle was kept throughout the experiment. Fecal pellets were collected using sterile tweezers at multiple timepoints and immediately stored at -80 °C. Offspring weight was measured at PND21. All experiments were conducted under approval of the Institutional Animal Care and Use Committee, UC San Diego IRB protocols S00227M and S18200. Collected fecal pellets were thawed and equally split into two aliquots: one for untargeted metabolomics and one for shotgun metagenomic sequencing. Samples for untargeted metabolomics were extracted using 10 μL of cold 50% (v/v) methanol (MeOH) per 1 mg of fecal pellet. Following the addition of the extraction solvent, samples were homogenized using a 5 mm stainless steel bead in a TissueLyser II (QIAGEN) for 5 min at 25 Hz and then incubated at 4 °C for 30 min. Samples were subsequently centrifuged at 21,130 *g* for 3 min, and supernatants were collected in a 96-well plate for untargeted metabolomics analysis.

### UHPLC-MS/MS Data Acquisition

Samples were randomized and analyzed using an untargeted metabolomics analysis platform comprising an UltiMate 3000 LC system (Thermo Fisher Scientific) coupled to a Q-Exactive Orbitrap mass spectrometer (Thermo Fisher Scientific). The chromatography system consisted of a Kinetex C18 column (Phenomenex) and a mobile phase of solvent A (water + 0.1% formic acid) and solvent B (acetonitrile + 0.1% formic acid). A representative linear gradient was run with a flow rate of 0.5 mL/min as follows: 0-1 min 5% B, 1-7 min 98% B, 7-7.5 min 98% B, 7.5-8 min 5% B, and 8-10 min 5% B. MS/MS data were acquired in data-dependent acquisition (DDA) mode using positive electrospray ionization (ESI+). The MS scan range was set to 100 - 1500 *m/z* with a resolution at *m/z* 200 set to 35,000 with 1 microscan. Automatic gain control (AGC) was set to 5E4 with a maximum injection time of 100 ms. Up to 5 MS/MS (TopN = 5) spectra per MS1 were collected with a resolution at *m/z* 200 set to 35,000 with 1 microscans and AGC target of 5E4. The isolation window was set to 3.0 *m/z* and normalized collision energy was set to a stepwise increase of 20, 30, and 40 eV and a dynamic exclusion of 10 s.

### UHPLC-MS/MS Data Processing

Acquired .raw data were converted into .mzML open-access format using ProteoWizard MSConvert (32) and deposited in GNPS/MassIVE under the accession numbers MSV000089558 and MSV000092652. Feature detection and extraction were performed via MZmine 4.1 (33) using batch processing. The .mzbatch files used for processing can be found on the associated GitHub page. Briefly, mass detection was performed and ions acquired between 0 and 10 min, with MS1 and MS2 noise levels set to 5E4 and 1E3 respectively, were retained. Chromatogram builder parameters were set at 5 minimum consecutive scans, 1E5 minimum absolute height, and 10 ppm for *m/z* tolerance. Smoothing was applied before local minimum resolver, which had the following parameters: chromatographic threshold 85%, minimum search range retention time 0.2 min, minimum ratio of peak top/edge 1.7. Then, 13C isotope filter and isotope finder were applied. Features were aligned using join aligner with weight for *m/z* set to 80 and retention time tolerance set to 0.2 min. Features not detected in at least 2 samples were removed before performing peak finder. Ion identity networking and metaCorrelate were performed before exporting the final feature table. The GNPS and SIRIUS export functions were used to generate the final feature tables containing peak areas and the .mgf files necessary for feature based molecular networking (FBMN) (34), performed in GNPS2 (35), and molecular class prediction via CANOPUS (36), performed in SIRIUS 6.1 (37). FBMN parameters were set as follows for both networking and library annotation: 0.02 for both precursor and fragment ion tolerances; 0.7 minimum cosine score, and 5 minimum matching peaks. The propagated candidate bile acid library (38) was used to generate putative annotations, which were validated for the presence of diagnostic MS/MS fragment ions using MassQL (39) bile acid specific queries for the different bile acid steroid core hydroxylated statuses (https://massqlpostmn.gnps2.org/). The following queries were used:

1. Monohydroxylated - QUERY scaninfo(MS2DATA) WHERE MS2PROD=341.28:TOLERANCEMZ=0.01:INTENSITYPERCENT=5 AND MS2PROD=323.27:TOLERANCEMZ=0.01:INTENSITYPERCENT=5
2. Dihydroxylated - QUERY scaninfo(MS2DATA) WHERE MS2PROD=339.27:TOLERANCEMZ=0.01:INTENSITYPERCENT=5 AND MS2PROD=321.26:TOLERANCEMZ=0.01:INTENSITYPERCENT=5
3. Trihydroxylated - QUERY scaninfo(MS2DATA) WHERE MS2PROD=337.25:TOLERANCEMZ=0.01:INTENSITYPERCENT=5 AND MS2PROD=319.24:TOLERANCEMZ=0.01:INTENSITYPERCENT=5
4. Tetrahydroxylated - QUERY scaninfo(MS2DATA) WHERE MS2PROD=335.24:TOLERANCEMZ=0.01:INTENSITYPERCENT=5 AND MS2PROD=317.23:TOLERANCEMZ=0.01:INTENSITYPERCENT=5
5. Pentahydroxylated - QUERY scaninfo(MS2DATA) WHERE MS2PROD=333.22:TOLERANCEMZ=0.01:INTENSITYPERCENT=5 AND MS2PROD=315.21:TOLERANCEMZ=0.01:INTENSITYPERCENT=5

Metabolic features were traced across the two cohorts using classical molecular networking with Min Cluster Size set to 0. Matching pairs were obtained after filtering the merged_pairs.tsv for difference in parent mass <; 0.02, difference in retention time (RT) <; 0.3 min, modified cosine score > 0.7, and removing matches within the same cohort. Networking jobs are available for download at the following links:

1. Antepartum cohort - https://gnps2.org/status?task=80df966e597544b5b798d8571e7a1352
2. Postpartum cohort - https://gnps2.org/status?task=d00e383149bf49a0b037ccbc399ab6c5
3. Cross-cohort network - https://gnps2.org/status?task=5b5f690d5c20443585f62c1183baa633

### UHPLC-MS/MS Data Analysis

Feature tables from untargeted metabolomics analyses were imported in R 4.2.2 (R Foundation for Statistical Computing, Vienna, Austria) for downstream data analyses. Data quality was checked by examining total extracted peak areas, sample internal standard (IS), and QCmix, a reference sample containing 6 standards and acquired every 10 samples throughout the run. Blank filtering was performed by removing features when mean peak areas across all samples were not at least 5 times the one observed in blanks. Principal component analysis (PCA) and partial least square discriminant analysis (PLS-DA) were performed after robust center log ratio transformation (rclr) via the package ‘vegan v 2.6’ and ‘mixOmics v 6.22’ (40, 41). PERMANOVA was used to evaluate group centroid separation after dimensionality reduction. Performances of the PLS-DA models were evaluated using leave-one-out (loo) cross-validation and 999 permutations. Models with classification error rate (CER) <; 0.5 are considered discriminatory. Variable importance (VIP) scores were extracted for each feature and features with VIPs > 1 were considered significant for discrimination. Upset plots were generated using the package ‘UpSetR v 1.4’ (42). The natural log ratios of the significant features extracted from the PLS-DA models, with AMP associated features at the numerator and PBS associated features at the denominator, were plotted over time and significance was tested via repeated Welch’s *t*-test followed by Benjamini-Hochberg (BH) correction. Metabolomics data was integrated with metagenomics data via DIABLO (43). The model was built on samples for which both sequencing and metabolomics data was available retaining the the top 20 most discriminant feature for each omics block. Performance was evaluated using loo cross-validation and correlations > 0.7 between the features were visualized via a circos plot.

### Bile Acid Annotation

To further investigate and annotate molecules of interest, one single fecal sample was reinjected multiple times in the Q-Exactive Orbitrap mass spectrometer system with different acquisition parameters. Data were both acquired in positive and negative ESI mode and with different resolutions (up to 140,000), AGC targets (up to 5E5), and in-source energies (0, 10, 50, 80 eV). Additionally, the same fecal sample was also analyzed on a Bruker timsTof Pro2 mass spectrometer coupled to an Agilent HPLC system. Chromatographic separation was performed on a reverse-phase HPLC C18 column (2.1 mm x 120 mm) using 0.1% formic acid in water (A) and 0.1% formic acid in acetonitrile (B) as the mobile phase. The chromatography ran on a 12 min gradient: 0-0.50 min 5% B, 1.10 min 25% B, 7.50 min 40% B, 8.50 min 99% B, 10.10 min 5% B, 12.00 min 5% B. The column compartment was kept at 40 °C and the sample injection volume was 3 μL with a flow rate of 0.5 mL per min. Detection was performed in positive ESI mode using a DDA method with TIMS-MS-PASEF with the following parameters: nebulizer gas pressure at 2.2 bar, dry temperature 220 °C, gas flow 10l/min. The mass range was collected from 20 *m/z* to 1300 *m/z*, the PASEF used a 100 ms ramp time and 100 ms accumulation time, and the 1/K_0_ ranges from 0.80 Vs/cm^2^ to 1.2Vs/cm^2^. The raw data file was then imported into Metaboscape 2025b, where it was processed as a project with positive polarity and the T-ReX 4D (LC-TIMS-QTOF) workflow. The filter parameters included minimum 1 feature for extraction and presence in a minimum of 1 analysis. The default workflow methods and calibration methods available were used to process the features from the sample. Theoretical reference collision cross-section (CCS) values were predicted in the CCS-Predict Pro 2025 model.

### Metagenomics Sequencing

Aliquots of the fecal samples were transferred to the UC San Diego Microbiome Core to perform DNA extraction as previously described (44). Briefly, samples were purified via the MagMAX Microbiome Ultra Nucleic Acid Isolation Kit (Thermo Fisher Scientific) using a KingFisher Flex robot (Thermo Fisher Scientific). Blanks and mock communities (Zymo Research Corporation) were included in the analysis for quality control. DNA was quantified via a PicoGreen fluorescence assay (Thermo Fisher Scientific), and metagenomic libraries were prepared using the KAPA HyperPlus kit (Roche Diagnostics) following the manufacturer’s instructions via a EpMotion automated liquid handler (Eppendorf). Shallow shotgun sequencing was performed on an Illumina NovaSeq 6000 platform with paired-end 150 bp cycles at the Institute for Genomic Medicine (IGM), UC San Diego.

### Metagenomics Data Processing and Analysis

Demultiplexed FASTQ files were imported in Qiita (Study ID # 15345) and processed using the default workflow for metagenomics data (45). Briefly, adapters and host genome (mouse) sequences were removed using qp-fastp-minimap2 2023.12 (46). Then, qp-woltka 2024.09 was used to generate operational genomic units (OGUs) (47), which taxonomy was derived via the Web of Life (WoL2) reference database (48). Finally, OGUs were filtered against Greengenes2 (gg/2024.09) (49). The ‘phyloseq v 1.42’ package was used to manipulate the microbiome data (50). The median sequencing depth of the sample was ∼2,000,000 reads and samples with less than 500,000 reads were discarded from downstream analysis. For alpha diversity analysis, OGUs table was rarefied to minimum number of reads observed in a sample before calculating the Shannon diversity index. Before beta diversity analysis and differential abundant analysis, rare microbial features detected in less than 10% of the samples or with relative abundance in any sample <; 0.0001% were excluded. The non-rarefied OGUs tables were robust center log ratio (rclr) transformed using the package ‘vegan v 2.6’ before PCA via ‘mixOmics v 6.22’ (40, 41). Centroid separation was evaluated using PERMANOVA. Differential abundance analysis was performed using ALDEx2 (51). Species were considered significantly different between groups if adjusted p values after BH correction were <; 0.05. The natural log ratios of features associated with AMP exposure (numerator) or PBS (denominator) were plotted against time, as described for the metabolomics analysis. Species directionality overlap across the two cohorts was investigated via an Upset plot.

## Data and Code availability

Code used for the analysis and to generate the figures presented in this manuscript is available on GitHub (https://github.com/simonezuffa/Manuscript_AMP_Perinatal). Untargeted metabolomics data is publicly available in GNPS/MassIVE under the following accession codes: MSV000089558 (Antepartum cohort) and MSV000092652 (Postpartum cohort). Metagenomics data is available in Qiita (Study ID # 15345) and in ENA under the accession number PRJEB90218.

## RESULTS

Dams (n=20) received intravenous injections of either 150 mg/kg of ampicillin (AMP) or phosphate-buffered saline (PBS) for two consecutive days. Treatments were administered either before the estimated delivery time (gestational day [GD]17 and GD18; Antepartum cohort) or after birth (postnatal day [PND]2 and PND3; Postpartum cohort). Fecal samples were collected one day prior to treatment, during treatment, and on PND7, PND14, and PND21 for untargeted metabolomics analysis via UPLC-MS/MS and shotgun metagenomic sequencing (**Fig. 1A**). Offspring body weight was recorded at weaning (PND21) and was significantly higher in mice indirectly exposed to AMP during the perinatal period (**Fig. 1B**).

**Fig 1.**
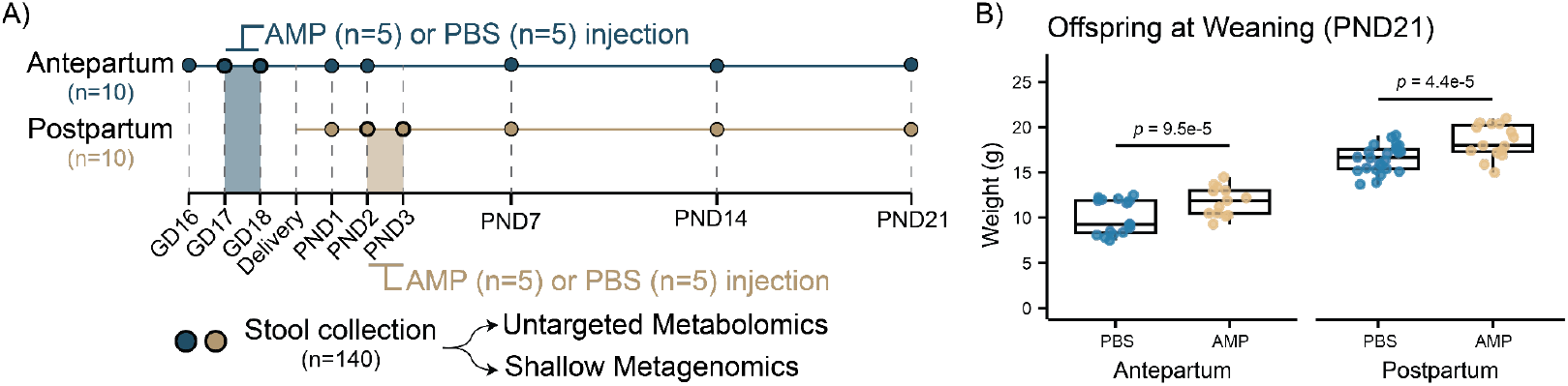
Perinatal maternal exposure to ampicillin increases offspring weight. **(A)** Two cohorts of dams (n=20) were injected intravenously with either AMP (150 mg/kg) or PBS for two days either antepartum (GD17 and GD18) or postpartum (PND2 and PND3). Fecal samples (n=131) were collected and analyzed via untargeted metabolomics and metagenomic sequencing. After quality control, 116 samples were included in the downstream analyses. **(B)** Offspring body weight was measured at weaning (PND21). Offspring of dams exposed to AMP had a significantly higher weight compared to the respective controls. *P* values were obtained using logistic regression adjusting for sex and litter. Boxplots show the first (lower) quartile, median, and third (upper) quartile. Abbreviations: AMP, ampicillin; GD, gestational day; PND, postnatal day; PBS, phosphate-buffered saline.

### Perinatal ampicillin exposure disrupts the maternal fecal microbiome

Metagenomic sequencing identified a time-dependent change in the maternal fecal microbiome throughout delivery and lactation (**Fig. S1A**) and alterations in response to AMP administration (PERMANOVA, *p* <; 0.001) in both the Antepartum and Postpartum cohorts (**Fig. 2A**). AMP significantly lowered Shannon alpha diversity (Wilcoxon test, *p* <; 0.05) during treatment (**Fig. 2B**), which then recovered at later timepoints (**Fig. S1B**). Differential abundance analysis using ALDEx2 identified 57 OGUs significantly altered by AMP in the Antepartum cohort (**Table S1**). Specifically, species belonging to the *Muribaculaceae* family, including *Muribaculum intestinale, Paramuribaculum intestinale*, and *Duncaniella dubosii*, and to the *Prevotella, Alloprevotella, Parasutterella*, and *Alistipes* genera were depleted, whereas species belonging to the *Enterococcus, Paenibacillus*, and *Staphylococcus* genera, such as *Enterococcus gallinarum, Paenibacillus cookii*, and *Staphylococcus xylosus*, were enriched in response to AMP (**Fig. 2C**). In contrast, 291 OGUs were altered in the Postpartum cohort (**Table S2**). AMP induced the enrichment of bacterial species belonging to the *Prevotella, Bacteroides, Escherichia, Phocaeicola, Alloprevotella*, and *Klebsiella* genera and the depletion of species belonging to the *Muribaculum, Duncaniella, Paramuribaculum, Blautia, Akkermansia* and *Roseburia* genera (**Fig. 2C**). Interestingly, looking at the directionality of significant overlapping OGUs between the two cohorts (n=19), only 17 showed a concordant reduction in response to AMP, whereas species-specific enrichments appeared to be cohort-dependent (**Fig. 2D**). Bacteria depleted in both cohorts encompassed short-chain fatty acid (SCFA) producing species belonging to the *Muribaculaceae* family, including *Muribaculum intestinale, Muribaculum gordoncarteri, Duncaniella dubosii, Paramuribaculum intestinale* and the genera *UBA7173, CAG-873*, and *CAG-485* (**Table S3**). Only two species were commonly enriched after AMP administration, which included *Edwardsiella ictaluri* and the *Bacteroidaceae* species *OM05-12 sp003438995*. Analysis of log ratios between AMP-enriched (numerator) or AMP-depleted (denominator) OGUs revealed that microbial disruption was transient and while it was restricted to the administration days in the Postpartum cohort, in the Antepartum cohort effects persisted until PND2 (**Fig. 2E**). Plotting the ratios of exclusively overlapping OGUs between the two cohorts, with concordant directionality (**Table S3**), recapitulated the observations obtained from the full model comparisons (**Fig. 2F**). Finally, differential pathway analysis in both cohorts showed concordant increases in butanoate metabolism, glycine, serine and threonine metabolism, sulfur metabolism, lysine degradation, pentose and glucuronate interconversions, and taurine and hypotaurine metabolism in response to AMP treatment (**Table S4**). Conversely, pathway depletion was cohort-dependent, with the Postpartum cohort showing decreased glycerolipid metabolism, mannose type O-glycan biosynthesis, carotenoid biosynthesis, and N-glycan biosynthesis.

**Fig 2.**
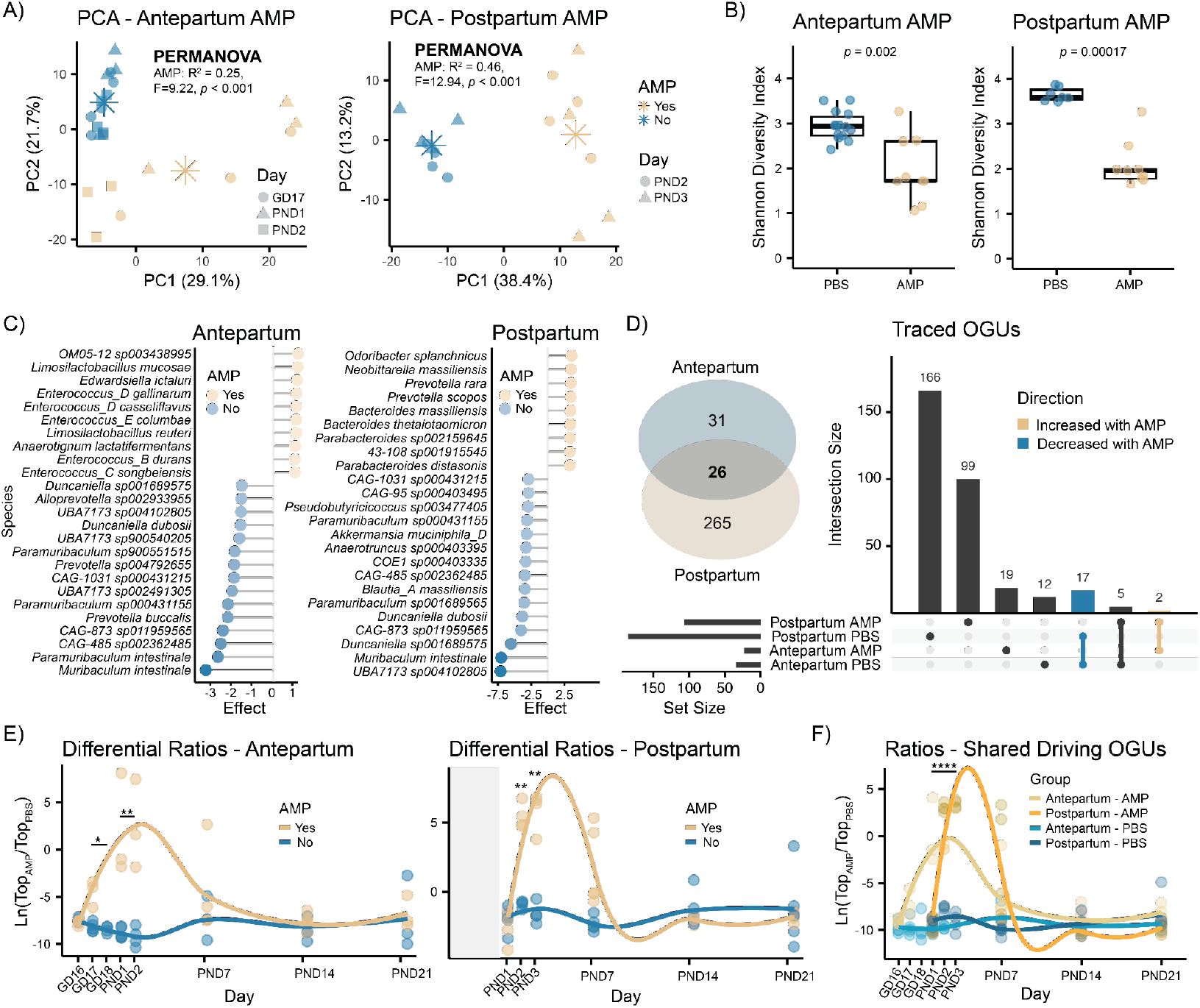
Perinatal ampicillin exposure alters the maternal fecal microbiome. **(A)** PCA of rclr transformed microbial features of fecal samples collected during treatment showed significant differences in response to AMP in both cohorts (PERMANOVA, *p* <; 0.001). **(B)** Shannon alpha diversity was significantly reduced in dams receiving AMP (Wilcoxon, *p* <; 0.05) followed by recovery at later timepoints (**Fig. S1**). **(C)** Top 10 enriched and top 15 depleted bacterial species in response to AMP, based on effect sizes obtained via ALDEx2. Only taxa with adjusted *p* <; 0.05 were included. **(D)** Overlapping bacterial species (n=26) altered in both cohorts following AMP treatment. Of these, 19 showed concordance in directionality, being either depleted (17) or enriched (2) in response to AMP. Commonly depleted taxa included species belonging to *Muribaculum, Paramuribaculum*, and *Duncaniella* genera. **(E)** Longitudinal log-ratio analysis of differentially abundant species identified via ALDEx2 per cohort. Features enriched in response to AMP were aggregated at the numerator, whereas depleted features were combined at the denominator. Significance was assessed using repeated Welch’s *t*-test with BH correction. **(F)** Log ratios of exclusively overlapping and directionally concordant species between cohorts across time. Ratio in the Postpartum cohort were normalized to the mean ratio difference between the two cohorts before treatment. Boxplots indicate the first (lower) quartile, median, and third (upper) quartile. Asterisks in PCA denote group centroids. Significance: * *p* <; 0.05, ** *p* <; 0.01, **** *p* <; 0.0001. Abbreviations: AMP, ampicillin; GD, gestational day; PND, postnatal day.

### Perinatal ampicillin exposure alters the maternal fecal metabolome

Untargeted metabolomics analysis of fecal samples showed AMP detection during administration and up to two days following the final injection (**Fig. 3A**). Unsupervised PCA revealed time-dependent shifts of the fecal metabolome (**Fig. S1C**) and clear separation (PERMANOVA, *p* <; 0.001) based on AMP exposure during treatment in both cohorts (**Fig. 3B**). Supervised PLS-DA models achieved near-perfect classification performance in both cohorts, highlighting the strong effect of AMP. Metabolic features driving group separation with VIP > 1 were extracted from both models and further investigated. A total of 4045 features were significantly altered in the Antepartum cohort in response to AMP (**Table S5**), while 2832 were altered in the Postpartum cohort (**Table S6**). The natural log ratios of the features increased (numerator) or decreased (denominator) after AMP exposure were analyzed in response to time (**Fig. 3C**). Interestingly, the effect of AMP was observable up to weaning day (PND21) in the Antepartum cohort, while the effect was restricted to the administration period only in the Postpartum cohort. To trace molecular features of interest across the two different cohorts, a cross-cohort molecular network with no clustering between the consensus MS/MS spectra was created. Network pairs were filtered to retain pairs with delta *m/z* <; 0.02, delta retention time <; 0.3 min, and cosine similarity > 0.7. A total of 668 features were matched across the two cohorts between the differential features of interest (**Table S7**). Notably, 560 of them showed the same directionality in both cohorts, with 302 decreased and 258 increased in response to AMP treatment (**Fig. 3D**). Feature annotation via FBMN and molecular class prediction via CANOPUS identified most of them as carnitines, bile acids, polyamines, and small peptides.

**Fig 3.**
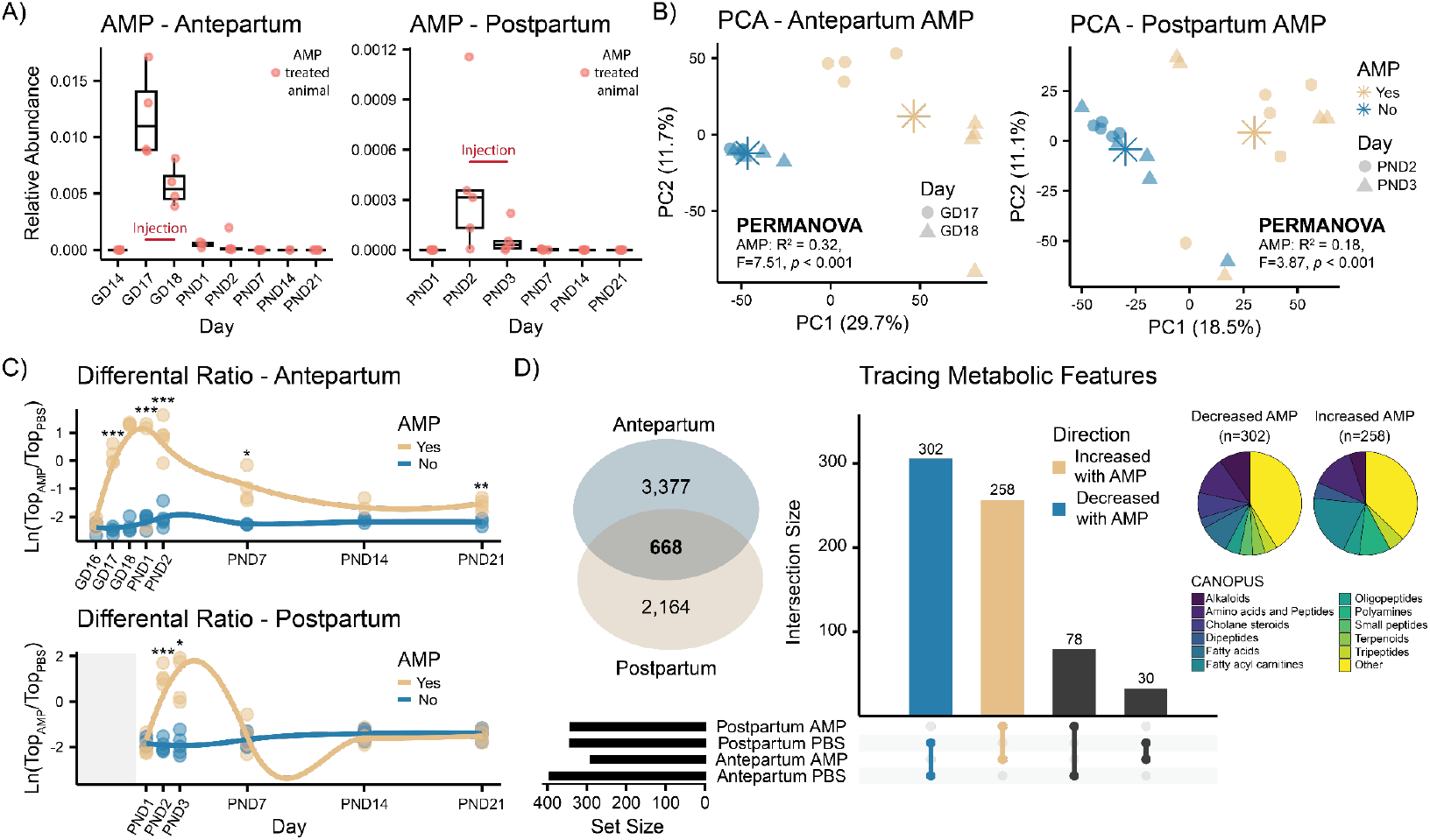
Ampicillin administration alters maternal fecal metabolic profiles. **(A)** Relative peak area abundance of detected fecal ampicillin ([M+H] *m/z* 350.1174, RT 2.7 min) in treated dams. No AMP was administered at the earliest timepoints (GD14 and PND1). **(B)** PCA of fecal samples collected during AMP exposure in the respective cohorts. Metabolic profiles were significantly altered based on exposure (PERMANOVA, *p* <; 0.001). Two outliers were observed amongAMP-treated animals in the Postpartum cohort. **(C)** Longitudinal analysis of the natural log ratios of features that were increased (numerator) or decreased (denominator) in response to AMP extracted from the PLS-DA models constructed using administration timepoints solely. Statistical significance was assessed using repeated Welch’s *t-*test with BH correction. **(D)** Overlapping metabolic features (n=668) altered in both cohorts following AMP treatment. Of these, 560 showed concordance in directionality, being either increased (n=258) or decreased (n=302) in response to AMP. Molecular classes of concordant molecular features were predicted using CANOPUS. Classifications are reported in the pie charts. Boxplots show first (lower) quartile, median, and third (upper) quartile. Asterisks in PCA represent group centroids. Significance: * *p* <; 0.05, ** *p* <; 0.01, *** *p* <; 0.001. Abbreviations: AMP, ampicillin; GD, gestational day; PND, postnatal day.

### Fecal acylcarnitines and bile acids are altered in response to ampicillin exposure

Several annotated and predicted acylcarnitines were altered in maternal feces following AMP exposure (**Fig. 4A**). In both cohorts, more than 40 distinct carnitines were consistently increased during antibiotic treatment. These included short and medium-chain carnitines, such as butyryl-carnitine, valeryl-carnitine, hexanoyl-carnitine, octanoyl-carnitine, decanoyl-carnitine, as well as carnitines with longer chain length and/or hydroxyl groups, such as hydroxy-tetradecanoyl-carnitine. Through information propagation in the submolecular networks, CANOPUS class prediction, and MassQL validation of the diagnostic fragment ions for carnitines (*m/z* 60.0813 and *m/z* 85.0287), more than 30 carnitines were also putatively annotated (**Table S8**). The bile acid composition was also affected by AMP administration. In both cohorts, 14 different di-, tri-, and tetra-hydroxylated bile acids, which included the putatively annotated deoxycholic acid, were altered in response to AMP (**Table S9**). The natural log-ratio of these bile acids alone recapitulated the longitudinal observations previously obtained using all differential metabolic features (**Fig. 4B**). We further investigated two unannotated bile acids (*m/z* 588.3740 and *m/z* 695.3790) that were consistently elevated in dams treated with AMP. These were two trihydroxylated bile acids, putatively cholic acid (CA) and taurocholic acid (TCA), given the presence of the respective diagnostic ions for the bile acid steroid core and taurine (**Fig. S2A**). Interestingly, they were both ammonium adducts [M+NH4]+, with the other ion forms being lower in abundance and failing to trigger MS/MS acquisition (**Fig. S2B**). SIRIUS chemical formula prediction for *m/z* 695.3790 returned C_32_H_55_NO_12_S, corresponding to a taurocholic acid (C_26_H_45_NO_7_S) conjugated to a hexose sugar (C_6_H_12_O_6_) via water loss (**Fig. S2C**). The spectrum was also further investigated using a timsTOF, where the [M+H]+ adduct form returned a ΔCCS% of 5.8% relative to a theoretical TCA-hexose conjugate. Querying these two spectra against the public metabolomics repositories via tissueMASST(52) revealed they were predominantly detected in the mouse feces and gastrointestinal tract tissues and more rarely in human samples (**Fig. 4C**). The unstructured search output obtained from tissueMASST showed that these spectra were also detected across several murine datasets lacking detailed metadata, but associated with antibiotics use. Suggesting that these bile acid derivatives may serve as markers of antibiotic exposure. Finally, multi-omics integration via DIABLO (43), highlighted that both *Muribaculum intestinale* and *Paramuribaculum intestinale* had a strong negative correlation to several medium to long-chain carnitines, that we have previously shown to be enriched in response to AMP (**Fig. 4D**).

**Fig 4.**
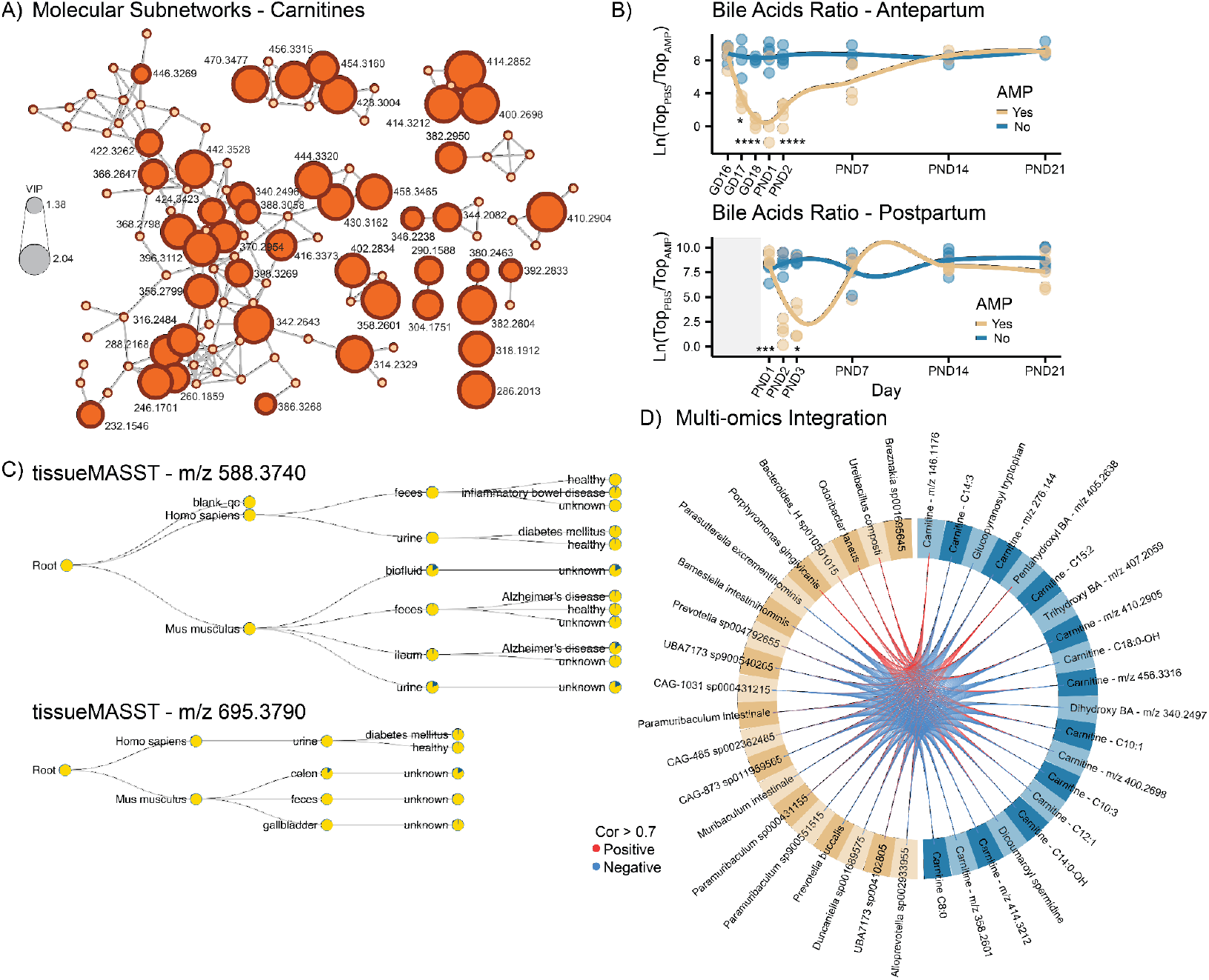
Fecal metabolic features altered by ampicillin treatment in dams. **(A)** Submolecular network of carnitines. Node size reflects the VIP scores from the PLS-DA classification. The majority of detected carnitines were enriched in response to AMP and putative annotations are provided in **Table S8**. (**B**) Longitudinal analysis of the natural log ratios of bile acids altered by AMP treatment across two cohorts. The effect was restricted to the treatment period in the Postpartum cohort, while it persisted until PND2 in the Antepartum cohort. Statistical analysis was performed using repeated Welch’s *t*-test followed by BH correction. Significance: * *p* <; 0.05, *** *p* <; 0.001, **** *p* <; 0.0001. (**C**) tissueMASST output from MS/MS searches. The molecules were predominantly detected in murine samples. (**D**) Circos plot obtained from multi-omics integration via DIABLO, combining metabolomics and metagenomics data in the Antepartum cohort. Red lines indicate positive correlations, and blue lines indicate negative correlations. Only correlations with coefficient > 0.7 are visualized. The model yielded a classification error rate (CER) of 0.16.

## DISCUSSION

The perinatal period, immediately before or after delivery, represents a critical window for the newborn’s long-term development. As maternal microorganisms and metabolites seed the infant at birth (53), disruptions to maternal gut microbial communities and their metabolic functions may adversely affect the growth and immune developmental trajectories of offspring. In this study, we explored the impact of ampicillin (AMP) on maternal fecal microbial and metabolic profiles. Notably, offspring indirectly exposed to AMP via maternal treatment exhibited increased weight at weaning (PND21) compared to unexposed controls. Direct exposure to broad-spectrum antibiotics during the first months of life is already known to increase infant weight at one year or later in life (13, 54). Supporting this, a recent large retrospective study of over 223,431 children further showed that GBS-IAP was associated with sustained increases in infant body mass index (BMI) from 6 months to 5 years of age, compared to no antibiotic exposure (55).

Across both cohorts, AMP treatment resulted in clear disruptions to maternal fecal microbiome composition. Interestingly, the proportion of overlapping, altered bacterial species was low, suggesting that many observed changes were cohort-dependent, potentially influenced by initial maternal colonization patterns or environmental variation. Nonetheless, several consistent findings emerged. We observed a reproducible depletion of species within the *Muribaculaceae* family, including *Muribaculum intestinale, Muribaculum gordoncarteri, Duncaniella dubosii, Paramuribaculum intestinale*, and the genera *UBA7173, CAG-873*, and *CAG-485*. Members of this family are known producers of SCFAs, which are beneficial to host health (56), and are involved in cross-feeding networks with other gut commensals such as *Bifidobacterium* and *Lactobacillus (57)*. Among these, *M. intestinale* has been shown to restrict pathogen colonization by converting succinate to propionate (58), a mechanism with potential relevance to early-life pathogen colonization resistance. Succinate itself is known to promote colonization by pathogens such as *Clostridium difficile (59)*, suggesting that *M. intestinale* could play a protective role. Additionally, *M. intestinale* can produce a proinflammatory cardiolipin (60), which may help prime the developing immune system. Similarly, *D. dubosii* has been associated with immunomodulatory effects via tryptophan metabolism (61). In the Antepartum cohort, AMP also reduced the abundance of several *Alistipes* species, including *A. communis* and *A. timonensis*, SCFAs producers with reported immunomodulatory properties (62). In the Postpartum cohort, *Akkermansia muciniphila*, a mucin-degrading bacterium with protective effects against metabolic disorders and diet-induced obesity (63), was likewise depleted. Early-life supplementation with *A. muciniphila* has been associated with increased intestinal goblet cell number, reduced adiposity, and improved glucose homeostasis in adulthood (64). The depletion of these beneficial taxa in the dams during the perinatal period may reduce their transmission to the gastrointestinal tract of offspring, impairing early-life colonization and immune system development, and potentially increasing susceptibility to pathogen colonization.

The ampicillin treatment also altered the maternal fecal metabolome. We observed elevated levels of multiple acylcarnitines, including several previously uncharacterized ones that we annotated via molecular networking. These molecules, important in energy metabolism via β-oxidation (65), have been reported to accumulate in the cecum, but not plasma, of antibiotic-treated mice (66). Acylcarnitines can also favor the growth of *Enterobacteriaceae* species (67), including opportunistic pathogens such as *Escherichia coli* and *Klebsiella* spp., both of which were enriched following AMP treatment in the Postpartum cohort. These organisms can also develop AMP resistance (68, 69) and they may be vertically transmitted to the infant during delivery or lactation. Similarly, in the Antepartum cohort, we observed enrichment of *Enterococcus* and *Paenibacillus* species, which can also acquire AMP resistance (70, 71).

Bile acid composition was another metabolic domain affected by AMP. These steroidal molecules, which are obtained from cholesterol metabolism, are closely related to gut microbiome activity, as gut bacteria modify primary bile acids through diverse enzymatic pathways (31, 38). We observed consistent depletion of microbially derived bile acids, including putative deoxycholic acid (DCA) and ketodeoxycholic acid (ketoDCA), consistent with previous reports (72). DCA, in particular, inhibits *Clostridium difficile* spore germination and vegetative growth (73, 74), and its depletion might facilitate pathogen colonization in the offspring. Across both cohorts, we also observed a sustained enrichment of two previously uncharacterized bile acid conjugates: cholic and taurocholic acid bound to a hexose sugar, likely glucose, fructose, galactose, or an isomer. The detection of these two bile acid conjugates appeared to be relatively rare across public data but was mostly observed along the gastrointestinal tract in both mice and humans, as shown by tissueMASST (52). Their potential microbial origin was supported by matches returned exclusively from microbeMASST (75), with no matches against the other domainMASSTs (76, 77). Although the biological roles of these saccharide-conjugated bile acids remains unclear and largely uninvestigated, they may reflect microbiome-dependent alterations in carbohydrate metabolism, a pathway tightly linked to obesity and insulin resistance (78). Excess monosaccharides have been associated with low-grade inflammation, while fructose promoted hepatic lipid accumulation (79), and galactose is known to activate immune cells (80).

## CONCLUSION

In conclusion, our findings demonstrate that perinatal AMP exposure perturbs key maternal microbial taxa and associated metabolic pathways, specifically those involved in bile acid modification and energy metabolism. These alterations are likely to contribute to the observed increase in offspring weaning weight, and underscore the need for careful evaluation of the timing, necessity, and downstream consequences of broad-spectrum antibiotic use during the perinatal window.

## LIMITATIONS

The impact of indirect ampicillin exposure via maternal milk on the offspring was not investigated, due to challenges in sample collection from offspring and a high rate of cannibalism. Maternal stress during the perinatal period appeared to exacerbate cannibalistic behaviour, highlighting the need for careful evaluations of methodologies for antibiotic administration during this crucial window. The collection of low biomass samples from the offspring also posed a challenge and appropriate negative controls should be rigorously implemented in the future studies of this nature. Although the sample size was limited, the pronounced effect of the ampicillin treatment, coupled with longitudinal sampling and reproducibility across the two separate cohorts, strengthens the generalizability of our findings. Untargeted metabolomics data were acquired in two separate batches, one year apart, which is suboptimal for direct comparisons. Nevertheless, our approach of first analyzing datasets independently and subsequently tracking molecular features of interest via high spectral similarity mitigates potential batch-related discrepancies. Metabolic feature annotations were obtained from MS/MS spectral matching against GNPS reference libraries, corresponding to level 2 annotation according to the Metabolomics Standard Initiative (MSI).

## ACKNOWLEDGMENTS

This work was supported by the Eunice Kennedy Shriver National Institute of Child Health & Human Development (NICHD) under grant number P50HD106463. We thank the UC San Diego Microbiome Core for their contributions to sequencing.

## Authors Contributions

S.Z. performed metabolomics and metagenomics analyses, generated figures, and wrote the manuscript. S.Z., S.P.T., I.M., Y.E.A., V.D., K.E.K., G.N., B.H. performed untargeted metabolomics experiments.

F.A., A.K., E.S., C.M.T. designed and conducted the animal study.

G.Y.L., V.N., P.C.D., F. A., and S.M.T. secured funding and/or provided supervision. All authors reviewed and approved the manuscript.

