## Supplementary Figures for "Influence of perinatal ampicillin exposure on maternal fecal microbial and metabolic profiles"

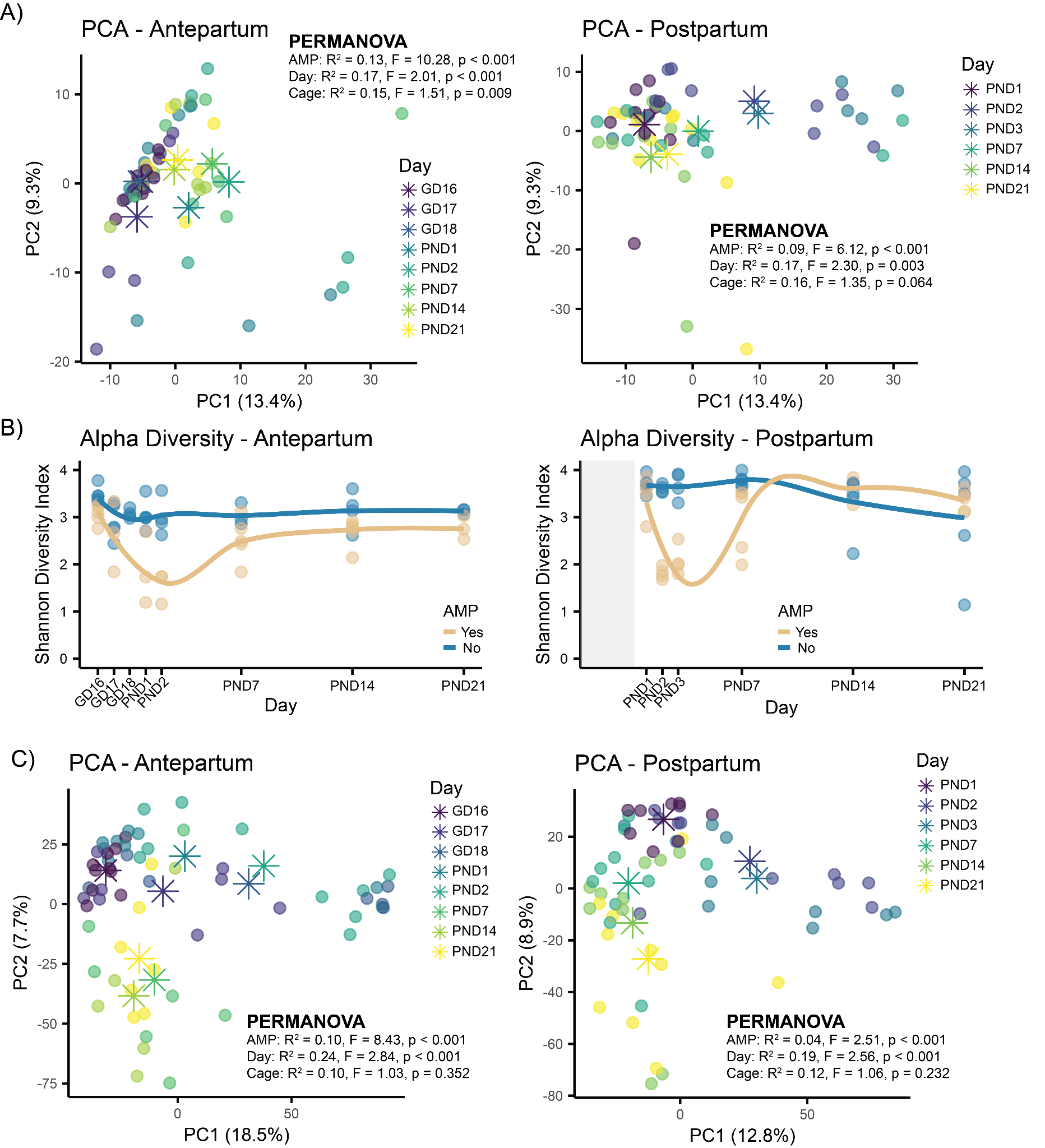

**Fig S1 Gut microbiome and metabolome time-dependent change**

**(A)** PCA of all fecal microbiome profiles collected either in the Antepartum or Postpartum cohort. In both datasets, a time-dependent change could be observed (PERMANOVA, *p* < 0.003) in addition to the AMP effect. A small inter-animal variance was also observed as the mice were single housed. Asterisks in PCA score plots represent group centroids. **(B)** Longitudinal analysis of Shannon diversity index identifies a reduced alpha diversity in correspondence of AMP treatment, which is then recovered at later timepoints. (**C**) PCA of all fecal metabolic profiles collected either in the Antepartum or Postpartum cohort. A time-dependent change of the metabolomes was observed (PERMANOVA, *p* < 0.001) in addition to the AMP effect. Abbreviations: AMP, ampicillin; GD, gestational day; PND, postnatal day.

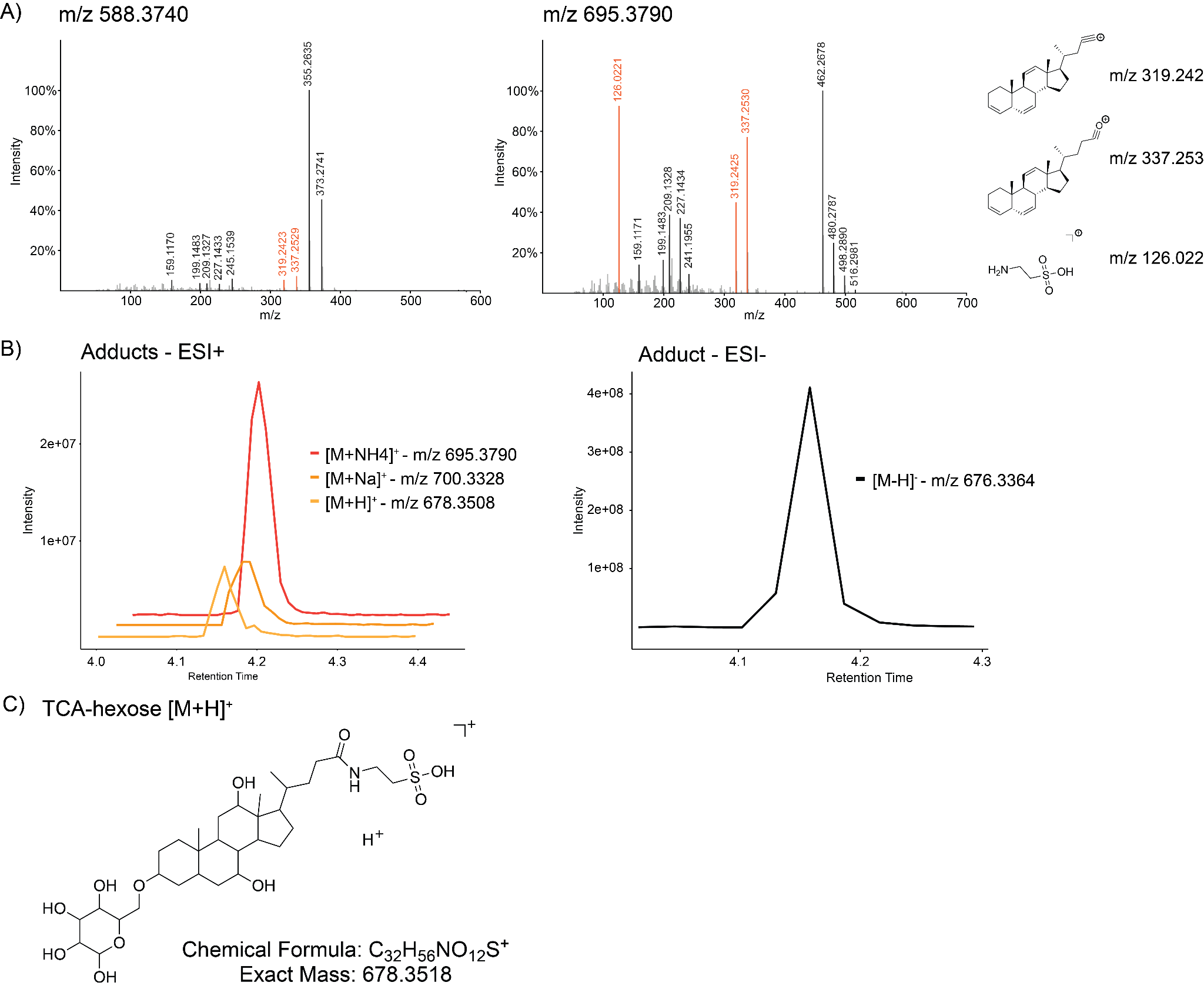

**Fig S2 Putative identification of uncharacterized bile acids spectra**

**(A)** MS/MS spectra of the two unknown bile acids. Given the presence of the diagnostic ions for trihydroxylated bile acids (*m/z* 319.242 and *m/z* 337.253) and taurine diagnostic ion (*m/z* 126.022) in one of the two, they were putatively annotated as conjugated forms of cholic and taurocholic acid respectively. **(B)** Extracted ion chromatograms for different ion adducts in relation to the *m/z* 695.3790 molecular feature. This feature of interest was an ammonium adduct, as both the proton and sodium adducts were detectable at the same retention time but they did not trigger MS/MS acquisition, possibly due to low abundance. (**C**) A putative structure of the taurine trihydroxylated bile acid conjugated to an hexose sugar.
